# Mosquito indolergic receptors belong to an ancient and functionally conserved Dipteran gene lineage

**DOI:** 10.1101/2020.08.04.236091

**Authors:** R. Jason Pitts, Shan Ju Shih, Jonathan D. Bohbot

## Abstract

Diptera is a megadiverse group of flies with sophisticated chemical detection systems, which exploits an incredible variety of ecological niches. Among the vast array of odorants in natural environments, indoles stand out as playing crucial roles in mediating fly behavior. In mosquitoes, indolic compounds are detected by an ancient class of conserved indolergic Odorant Receptors (*indolORs*). In this study, we have identified a set of 92 putative *indolOR* genes encoded in the genomes of Nematoceran and Brachyceran flies, resolved their phylogenetic relationships, and defined conserved elements in their gene structures. Further, we have quantified *indolOR* transcript abundance in the antennae of the housefly, *Musca domestica*, and have characterized MdomOR30a as a skatole receptor using a heterologous expression system. The presence of *indolORs* in species operating in different ecological contexts suggests that indoles act as pleiotropic signals for resource exploitation at multiple developmental stages. Further characterization of *indolORs* will impact our understanding of insect chemical ecology and will provide targets for the development of novel odor-based tools that can be integrated into existing vector surveillance and control programs.

## Introduction

Animals detect, track and locate a variety of resources in rapidly changing environmental conditions. Chemical cues are amongst the most predominant ecological drivers of animal behavior and exert major influences on resource acquisition [1, 2]. Perhaps nowhere in animal lineages is this dependency on olfactory information more evident than in insects, where accurate and timely chemical detection is necessary for life history traits such as foraging, toxin and predator avoidance, egg deposition, and mate selection. Indeed, insects, like other animals, possess highly evolved chemosensory systems that have allowed this large and diverse taxonomic group to exploit a dazzling array of ecological niches [3–8].

Despite this tremendous diversity, some interesting convergent principles may underlie animal olfactory-mediated behaviors. For example, highly ubiquitous compounds like indoles, which are produced by plants and microorganisms, can either attract or repel insects [9] and can evoke either pleasant or repugnant responses in humans [10, 11]. The behavioral valency for insects and the perceptual quality for humans should depend on the quantity of indolic compounds emitted from specific sources. While the molecular basis and ecological significance of indole sensing by humans is unknown, the genomes of mosquitoes encode a small number of highly selective and exquisitely sensitive *<u>indol</u>ergic <u>O</u>dorant <u>R</u>eceptors* (*indolORs*). *IndolORs* are remarkably conserved in mosquitoes [12–14], considering the highly dynamic nature of this gene family [12] and the ancient origin of the Anophelinae-Culicinae split that occurred 145-226 million years ago [15–17].

It is unclear whether indoles act as oviposition cues or as animal-host indicators [18, 19]. Furthermore, these heterocyclic compounds contribute to floral scent [20, 21] and are preferred by *A. aegypti* in the context of plant-host attraction [22]. The complex developmental expression patterns [12] and evolution of mosquito indolORs [13, 23, 24], in terrestrial adults and aquatic larvae, suggest that these two compounds play key roles in regulating inter-specific interactions in addition to their stage-specific functions. Mosquito species belonging to the subfamilies, Anophelinae and Culicinae, express both an indole-selective (*OR2*) and a skatole-selective (*OR10*) receptor. In Culicine mosquitoes, including members of the genera *Aedes*, a third paralog named *OR9* operates as a supersensitive skatole receptor during the aquatic stage [24]. The expression of *indolORs* in aquatic larvae [12] and in terrestrial blood- and nectar-feeding adults of different sex and species [23, 25] suggests that indolic compounds may have the intriguing property of influencing mosquito behaviors in multiple ecological contexts, including larval foraging, animal- and plant-host seeking, and oviposition site selection [19].

This paradigm may have important implications for olfactory coding whereby mosquitoes and other insects rely on various odor blends composed of combinations of commonly-occurring chemical signals, such as indoles, plus resource-specific odorants. It is therefore of interest to seek additional evidence in support of the proposition that other flies utilize parsimonious indole response mechanisms to optimize olfactory-mediated resource acquisition. Such findings would inform our understanding about the natural principles that have influenced the evolution of insect odor coding and behavior.

## Results

### The dipteran indolOR gene family is phylogenetically ancient

*IndolORs* are encoded by three genes in the yellow fever mosquito, *Ae. aegypti* (*ORs 2*, *9*, *10*). We used the conceptual translations of these genes in homology-based searches to explore the conservation of *indolORs* within the Diptera, focusing on species of medical and veterinary relevance in the families Culicidae, Glossinidae, Muscidae, and Psychodidae. The primary amino acid sequences of indolORs were used to build phylogenetic trees based on the Maximum-likelihood method (Fig. 1). Our analysis reveals a clear distinction in the evolutionary histories of *indolORs* between the lower flies (Nematocera) and higher flies (Brachycera), which may reflect novelties in their chemoreceptive functionalities (Fig. 1).

**Figure 1.**
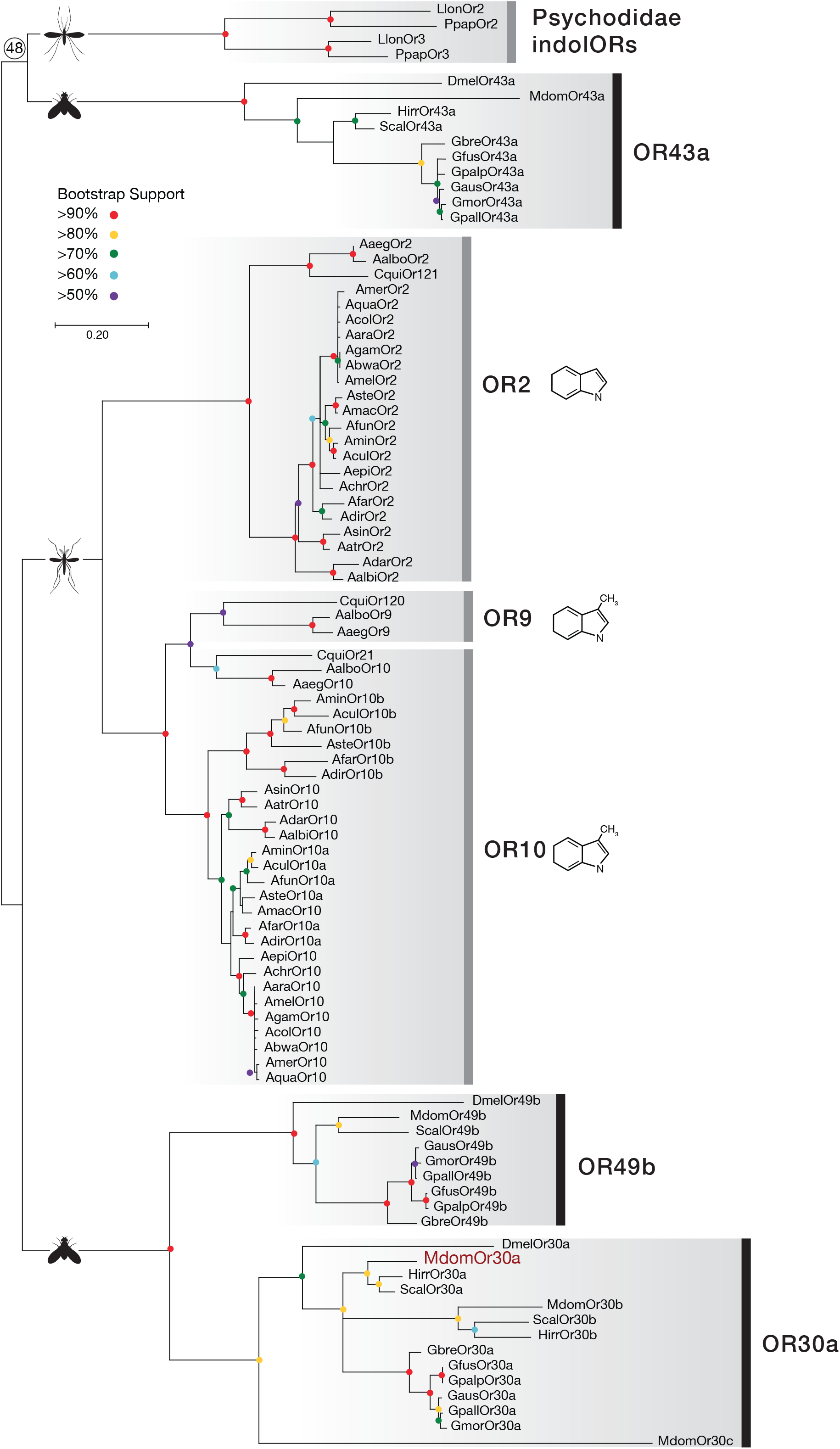
Dipteran indole-sensitive odorant receptor proteins (IndolORs) are phylogenetically ancient. Maximum Likelihood analysis of 92 fly indolORs homologs (Bootstrap support from 1000 replications color coded as indicated by the legend). Nematocera and Brachycera indolOR subclades are indicated by vertical grey and black bars, respectively. The bootstrap value supporting the Psychodidae and OR43a clades is shown above the split. Scale bar represents 0.2 changes per residue. For a full listing of indolORs and abbreviations, see Table S1.

Each of the *indolORs* exist almost exclusively as single-copy genes within the genomes we examined (Fig. 1). Subtle differences in the number of *indolORs* across genomes suggest single gene losses or gains. For example, the Anophelines generally encode two *indolOR* homologs, *ORs 2* and *10*, with species in the Cellia subgenus encoding a second *OR10* paralog, which we have named *indolOR10b* (Fig. 1). The Culicines encode three *indolORs*, including a distinct *OR9* group (Fig. 1). The genomes of *Lutzomyia* and *Phlebotomus* sandflies from the Psychodidae family, one of the oldest families of Diptera, also encode two *indolORs* [30, 31]. Most of the Brachyceran species encode exactly three candidate homolog *indolORs*, including single representative genes for *OR30a*, *OR43a*, and *OR49b*. We identified two of three homologs in *H. irritans*, with *OR49* apparently absent from the genome. Interestingly, we identified an additional paralog of *OR30a* in the genomes of *S. calcitrans* and *H. irritans*, which we have named *OR30b*, plus two additional paralogs in *M. domestica*, named *OR30b* and *OR30c* (Fig. 1). *OR30a* and *OR49b* clades are completely resolved with respect to the nematoceran lineages. Their branching pattern suggests that these genes evolved more recently than the OR43a clade as indicated by a potential common ancestor with the phylogenetically ancient Psychodidae *OR2* and *OR3* genes. Indeed, the branch leading to the Brachyceran OR43a clade indicates substantial amino-acid divergence suggesting a common ancestor with sandflies.

### Dipteran indolOR homologs are highly divergent

We examined the degree of homology across Diptera by aligning the amino acids sequences of a selected subset of receptors from *A. aegypti*, *P. papatasi*, *L. longipalpis*, *D. melanogaster*, and *M. domestica* (Fig. 2A). Amino acid conservation increases from the N-to C-terminus across all *indolORs* and is highest at the extreme C-terminal end of the receptors, encoded by their final exons. This C-terminal conservation is a general feature of insect odorant receptors [14]. Overall amino acid identities are typically between 30-40%, even when comparing different *indolORs* and across lower and higher flies (Fig. 2B). Highest identities of 50-80% are observed for intraspecies *indolORs* and for some intrafamily orthologs. For example, *A. aegypti* OR9 and OR10 are 70% identical, while *P. paptasi* OR3 and *L.longipalpis* OR3 are 81% identical (Fig. 2B). Another illustration of conservation that likely reflects their evolutionary histories is the relationships between the *M. domestica* and *D. melanogaster indolORs*. The OR30a, OR43a, and OR49b orthologs from these two species share 58%, 45%, and 63% amino acid identities, respectively (Fig. 2B). However, *Mdom*OR30a shares only 31% amino acid identity with *Dmel*OR43a, but 46% amino acid identity with *Dmel*OR49b, supporting a more distant evolutionary relationship between the *OR30a* and *OR43a* clades.

**Figure 2.**
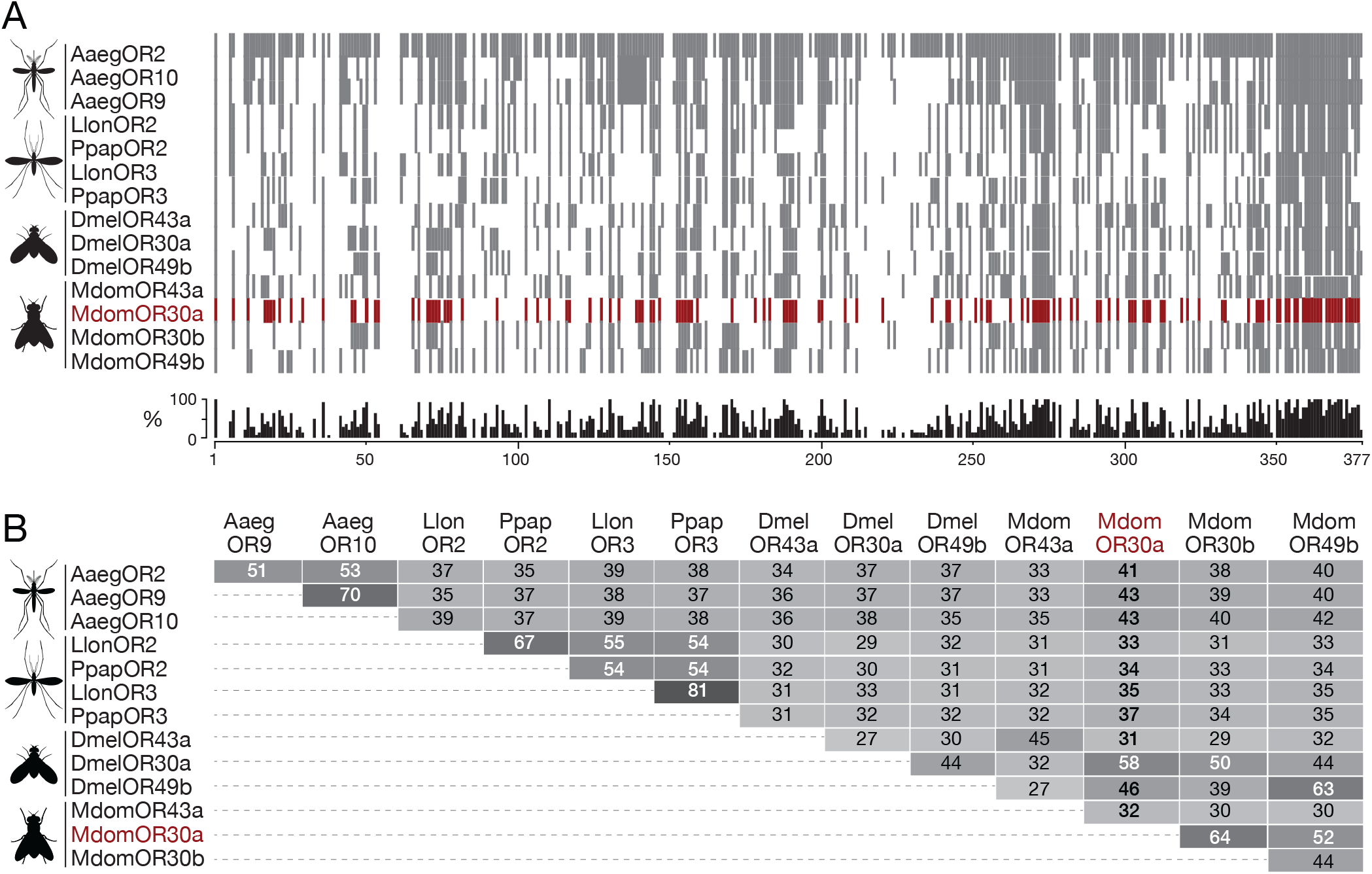
Dipteran indole-sensitive odorant receptors (IndolORs) proteins are highly divergent. A) Amino-acid sequence identity (grey) of indolORs from *Aedes aegypti* (Aaeg), *Lutzomyia longipalpis* (Llon), *Phlebotomus papatasi* (Ppap), *Drosophila melanogaster* (Dmel) and *Musca domestica* (Mdom) indolORs (OR2, OR9, OR10, OR3, OR43a, OR30a & OR49b). Percentage of amino-acid sequence identity is shown below. B) Amino-acid sequence identity matrix. Intensity of shading indicates magnitudes of homologies.

### Gene structure and microsynteny of Dipteran indolORs is conserved

Patterns of intron conservation (Fig. 3) and chromosomal synteny (Fig. 4) further support their evolutionary relationships. For example, the preservation of ancestral introns (A1-A6) indicate a common gene lineage for *indolORs*, while nematoceran-specific introns (N1, N2) indicate divergent evolution between lower and higher flies (Fig. 3). A2 is absent from the Culicidae lineage. Clustering of the OR2, 9, and10 genes, as well as conservation of neighboring orthologs in Culicidae indicated preservation of chromosomal segments in nematoceran (Fig. 4). Although the brachyceran *indolORs 30a*, *43a*, and *49b* are dispersed throughout their respective genomes, examples of preservation of orthologs are readily identified, especially for *indolOR49b* across the Drosophilidae, Muscidae, and Glossinidae families (Fig. 4).

**Figure 3.**
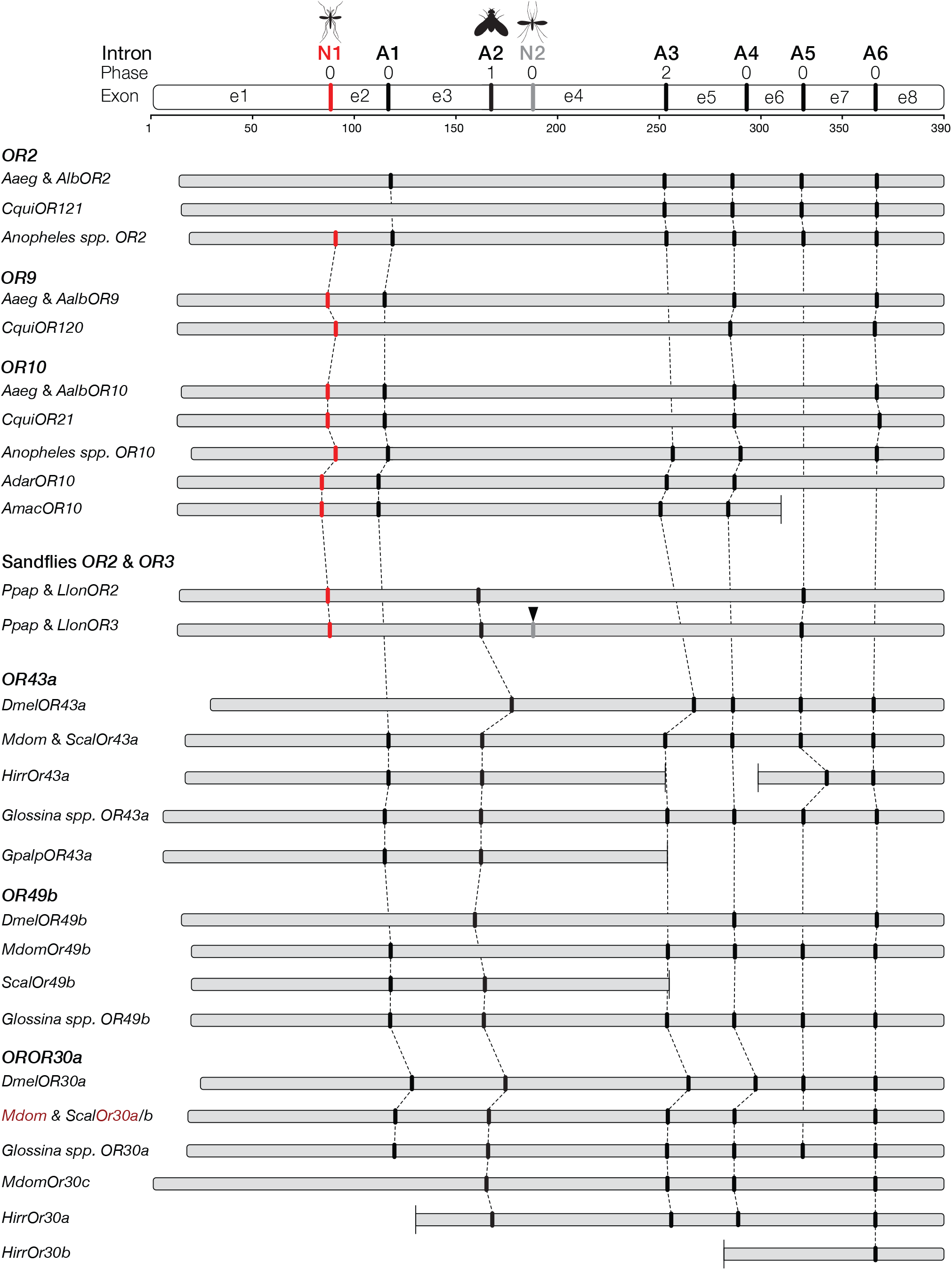
Dipteran indole-sensitive odorant receptor genes (*IndolORs*) exhibit conserved intron-exon structure. Exons (e1-8), positions and phases of ancestral (A1-6) and nematoceran-specific (N1, N2) introns in fly *indolOR* genes.

**Figure 4.**
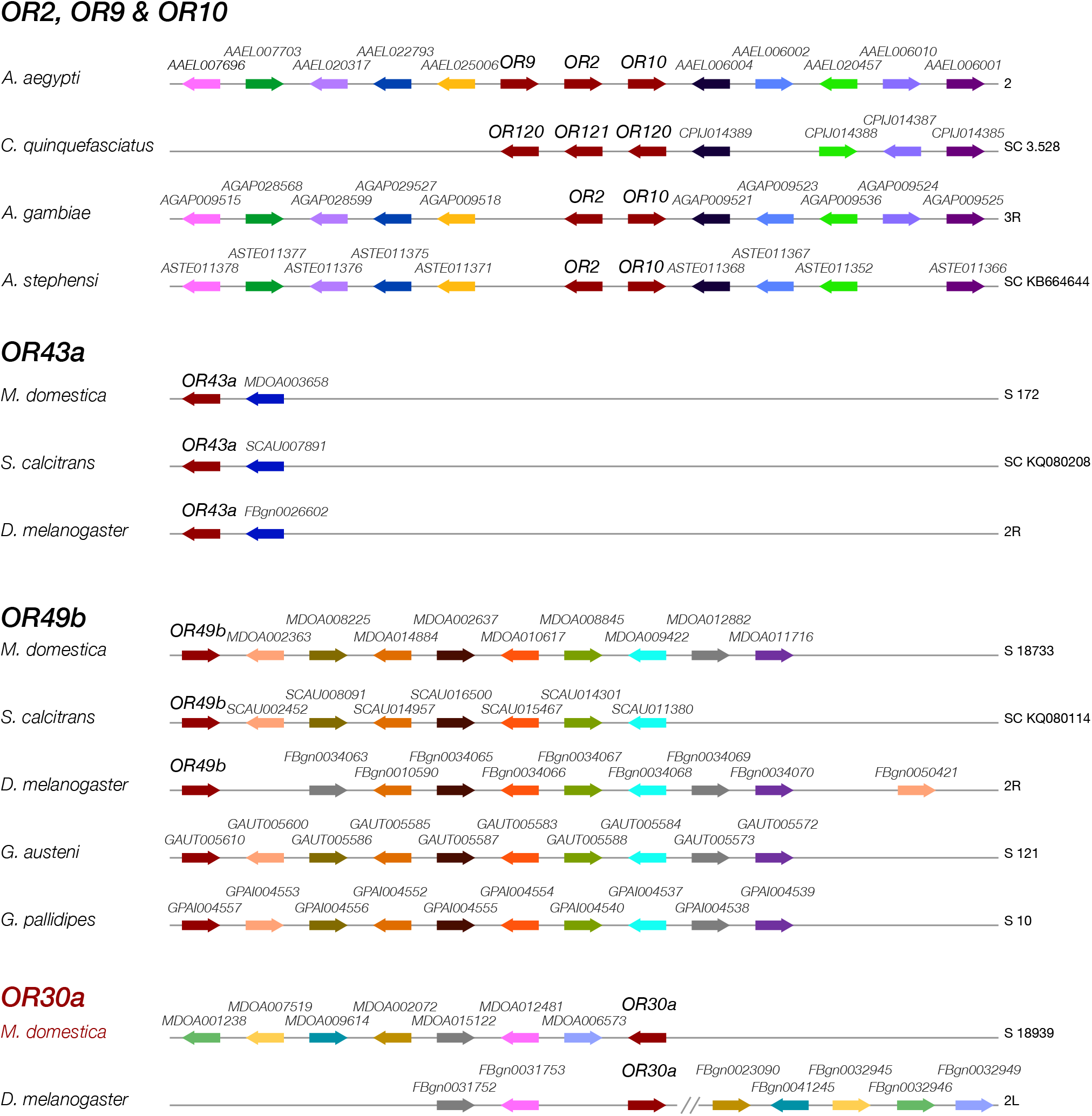
Microsyntenic relationships of dipteran indole-sensitive odorant receptor genes (*IndolORs*). Relative positions of IndolORs (red) and conserved neighboring genes (each homolog groups are color-coded) on supercontigs (SC), scaffold (S) and chromosomes. Direction of transcription is indicated by arrows. The color coding only applies within OR groups. Distances between genes are not drawn to scale. For a full listing of genes and abbreviations, see Supplemental Table 2.

### Musca domestica indolORs are expressed in the adult antennae

To examine the functional expression of brachyceran *indolORs*, we chose to focus on *Musca domestica*, a major global pest species that serves as a mechanical vector for bacterial pathogens, especially species in the genus, *Shigella*, that cause dysentery [32, 33]. We conducted an RNA sequencing analysis of antennae to assess transcript abundance as proxy for receptor expression. *IndolORs* were detected at levels suggesting that they are functional in this appendage in both sexes (Fig. 5). Abundances for all other transcripts, including additional chemoreceptors, are provided in Supplemental data. Among the antennal-expressed *indolORs* in *M. domestica*, OR30a displayed the highest abundance with an average TPM of 18.2 across both sexes, followed closely by *OR43a* with an average TPM of 17.7 (Fig. 5). The abundances of *OR30b* and *49b* were about 3 to 5-fold lower, which could reflect reduced expression in individual ORNs, the cumulative number of ORNs expressing each receptor, or both.

**Figure 5.**
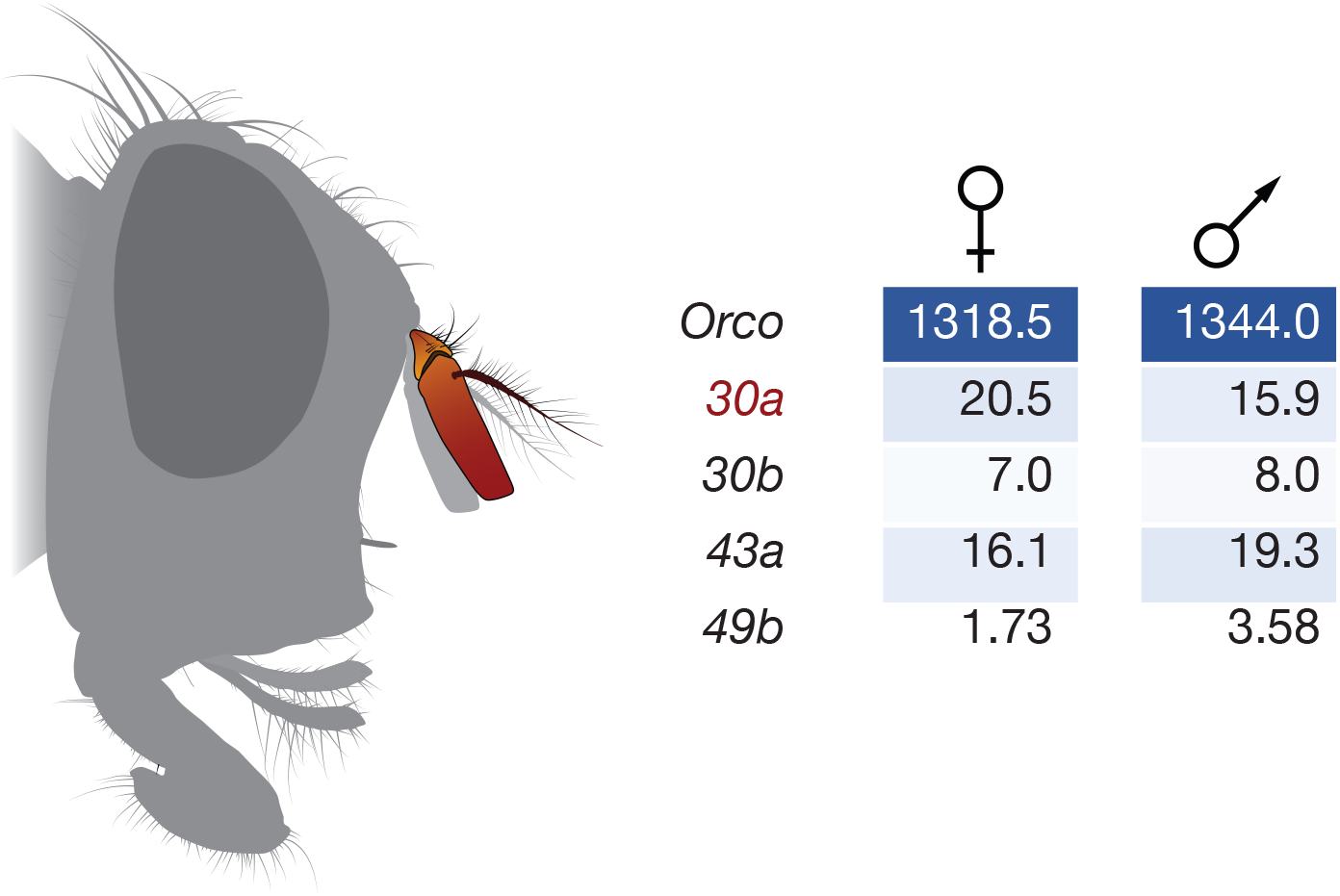
*M. domestica IndolORs* are expressed in adult male and female antennae. A) IndolOR transcripts, *30a*, *30b*, *43a*, and *49b*, and *Orco* are expressed in the antennae of male and female housefly. Numbers represent Transcripts Per Million (TPM).

### MdomOR43a is a skatole receptor

To test the potential functional orthology with mosquito indolORs, we utilized *Xenopus laevis* oocytes as a heterologous expression platform and the two-electrode voltage clamping technique to record the responses of MdomOR30a to a panel of structurally related indolic compounds. When co-expressed with MdomORco, MdomOR30a was highly selective towards skatole, followed by 2,3-dimethylindole and indole with a kurtosis value of 7.05 (Fig. 6A). MdomOR30a also displayed concentration-dependency of indolic responses over several orders of magnitude (Fig. 6B). With a half maximal effective concentration (EC_50_) in the low-mid nanomolar range, skatole was the most potent agonist odorant and was an order of magnitude lower than sensitivities to indole or 2,3-dimethylindole (Fig. 6C). This high degree of sensitivity to skatole is similar to those that have been reported for nematoceran, specifically mosquito, indolORs (Table 1).

**Table 1.**
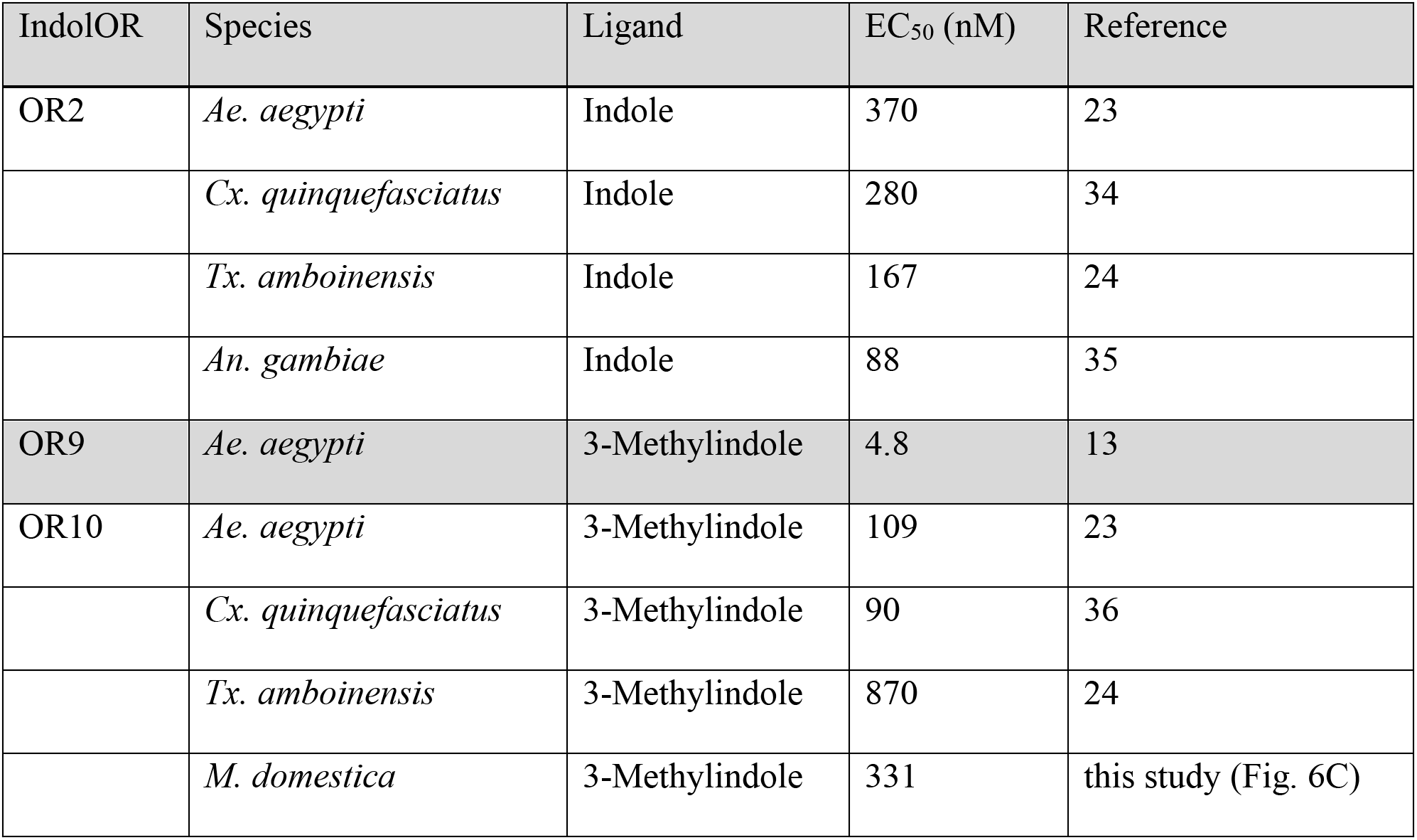
List of cognate pheromone and kairomone receptors deorphanized in the Xenopus laevis expression system.

| IndolOR | Species | Ligand | EC<sub>50</sub> (nM) | Reference |
| --- | --- | --- | --- | --- |
| OR2 | *Ae. aegypti* | Indole | 370 | 23 |
|  | *Cx. quinquefasciatus* | Indole | 280 | 34 |
|  | *Tx. amboinensis* | Indole | 167 | 24 |
|  | *An. gambiae* | Indole | 88 | 35 |
| OR9 | *Ae. aegypti* | 3-Methylindole | 4.8 | 13 |
| OR10 | *Ae. aegypti* | 3-Methylindole | 109 | 23 |
|  | *Cx. quinquefasciatus* | 3-Methylindole | 90 | 36 |
|  | *Tx. amboinensis* | 3-Methylindole | 870 | 24 |
|  | *M. domestica* | 3-Methylindole | 331 | this study (Fig. 6C) |

**Figure 6.**
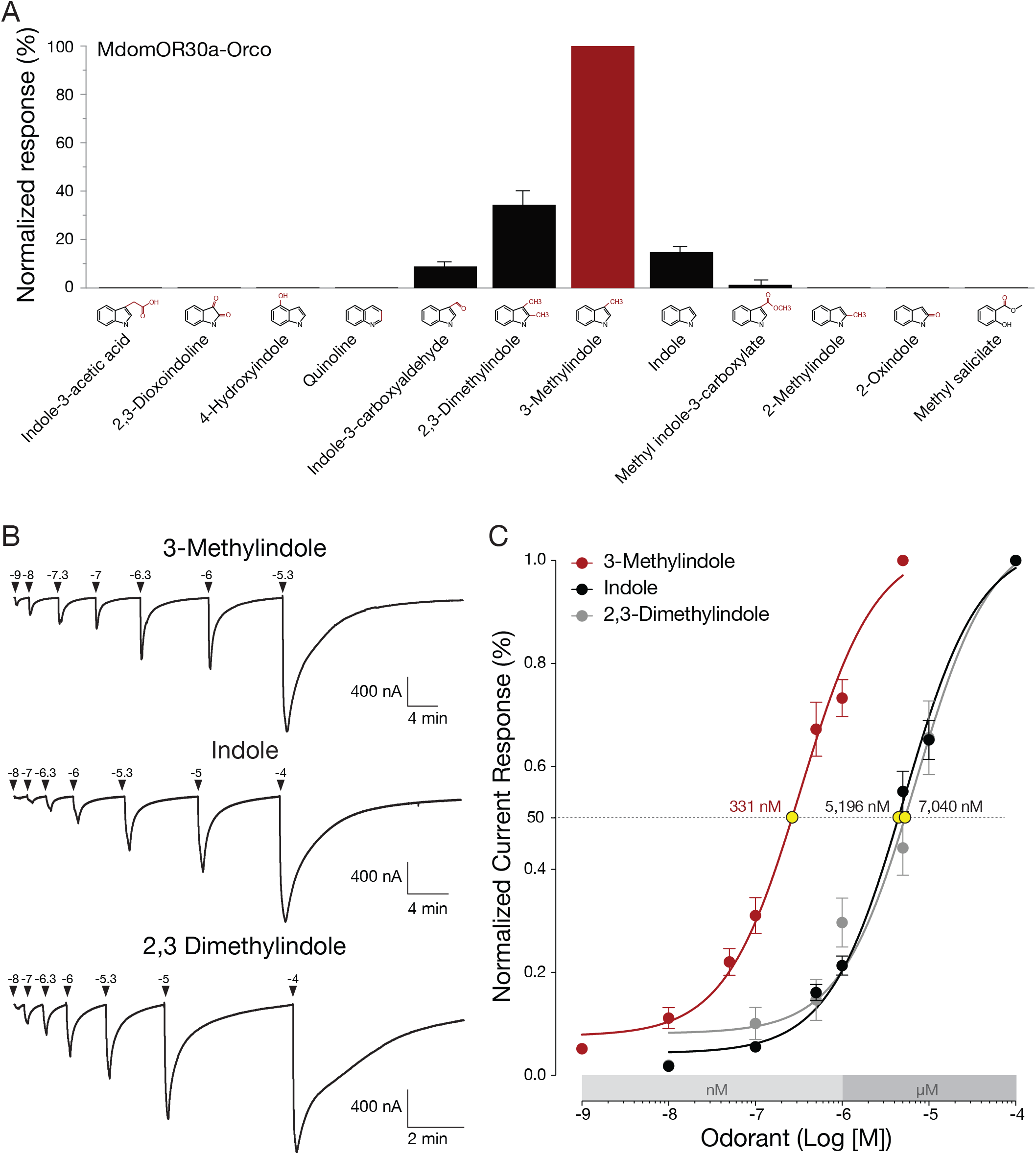
*M. domestica* OR30a is a skatole receptor. A) MdomOR30a is a 3-methylindole (skatole) selective receptor. Mean normalized oocyte responses using a panel of indolic compounds (n=10). B) Representative current traces elicited by 3-methylindole, indole and 2,3 dimethylindole. Arrowheads above traces indicate stimulus onset, while numbers indicate log molar concentrations of ligands. C) MdomOR30a responds to 3-methylindole in the nanomolar range (EC_50_ = 331 nM; n = 7). Both indole (EC_50_ = 5,196 nM; n = 10) and 2,3 dimethylindole (EC_50_ = 7,040 nM; n = 7) are less potent ligands, activating MdomOR30a in the high-nanomolar to low-micromolar range.

## Discussion

In this study, we propose that the brachyceran OR30a is an indolOR by virtue of its ligand selectivity, nanomolar sensitivity towards 3-methylindole, and gene structure. Until now, *IndolORs* were considered to be a Culicidae-specific expansion due to their ligand-binding characteristics [13, 24, 34–40] and amino-acid sequence conservation (50-70%) [12]. Candidate indolORs outside the Culicidae family display an amino-acid sequence identity significantly lower than within mosquitoes. Relaxing this qualifying criterion allowed us to identify putative *indolOR* homologs within the phylogenetically ancient Psychodidae family and more significantly in more distantly-related flies. OR30a, OR43a and OR49b form distinct and brachyceran-specific clades as reflected by their low amino-acid conservation (34-43%) with their mosquito counterparts and diverse syntenic relationships (Figs. 2, 4)

*IndolORs* are expressed in fly species that exhibit a great diversity of diets, oviposition habitats, and chemical ecologies. For example, mosquitoes, sand flies, stable flies, horn flies and tsetse flies are all hematophagous as adults. In the mosquitoes and sandflies, only the females bloodfeed while in the latter three groups, both sexes bloodfeed. Some of the brachyceran flies studied here, with the exceptions of *D. melanogaster* and Glossina species, produce maggots that feed on decaying matter (e.g., manure). The larvae of tsetse flies spend nearly all of their larval development within the female uterus, feeding on specialized glands.

*Drosophila melanogaster* also has 3 three putative *indolORs*, whose function are currently unknown. While OR43a and OR49b are expressed in the adult antennae [41], OR30a is only expressed during the larval stage [41–43], suggesting distinct developmental roles for indole detection. Little is known about the role of indoles in the chemical ecology of these insects. However, indole and 3-methylindole have been identified as potential food and oviposition attractants [44–46]. Expanding our analysis to additional dipteran genomes will likely uncover additional, divergent indolORs, thereby providing an opportunity to explore the roles of indoles in these species.

Female *M. domestica* flies are attracted by fetid scents signaling suitable oviposition sites, which are mainly provided by animal manure [47]. Similar odors are released by Sapromyiophilous plants to attract flies for pollination purposes [20]. In both cases, the attraction of *M. domestica* to oviposition and feeding sites is in part mediated by indoles (indole and 3-methylindole) sensitive antennal receptors in females as well as in males [47, 48] (Fig. 7). The discovery of a 3-methylindole sensitive and selective OR30a that is expressed in the antennae of the housefly, *Musca domestica*, provides a candidate molecular mechanism for the observed attraction of this insect to 3-methylindole [45, 49, 50]. Further experiments, such as gene knockout, will be required to establish a causal link between *OR30a* and 3-methylindole-mediated olfactory behaviors in flies. In addition, cellular-level expression studies and single-sensillum recordings are needed to resolve global antennal patterns of sensillar expression and the relationships between indolOR-expressing OSNs and their physiological functions in peripheral odor signaling.

**Figure 7.**
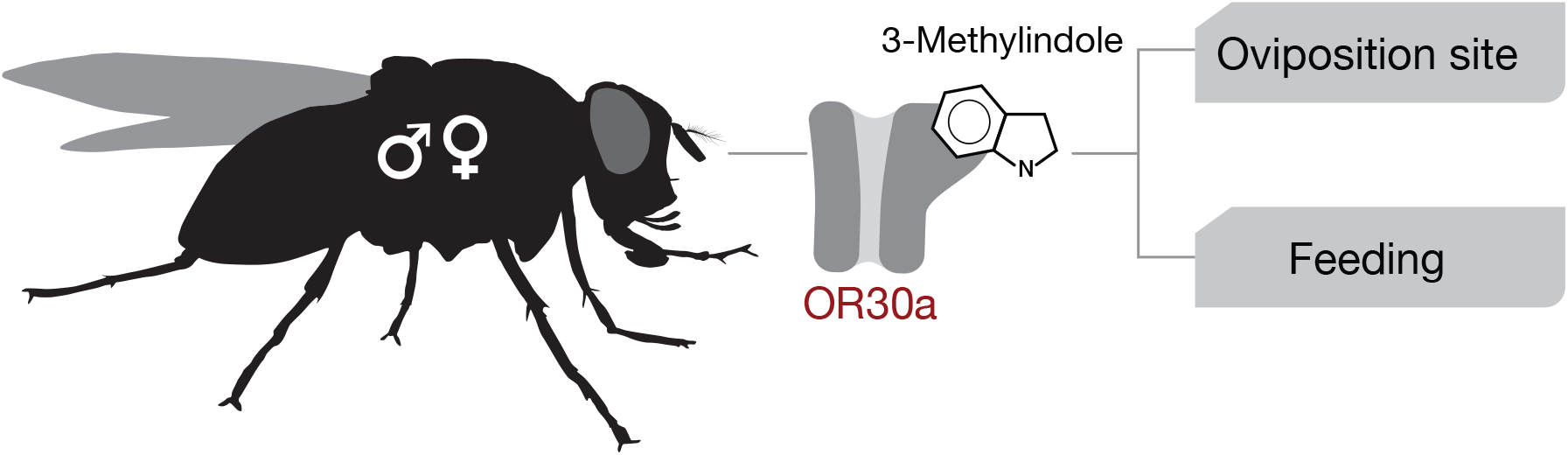
The MdomOR30a-3methylindole cognate pair mediates oviposition selection and food seeking behaviors. The expression of MdomOR30a in male and female houseflies supports a dual role of 3-methylindole in the attraction of flies to oviposition sites and food sources.

## Conclusions

In this study, we provide genetic, evolutionary and functional evidence supporting the hypothesis that *M. domestica* OR30a is a selective indolOR. The reconstitution of a sensitive and selective 3-methylindole receptor, recapitulating previous physiological studies [47, 48], paves the way for exploring the roles of indolORs to indole-sensing in insects exhibiting a broad variety of feeding and oviposition habits. Finally, the identification of *indolORs* across multiple dipteran lineages illustrates the multiple roles of indoles in insect chemical ecology and underscores their importance in olfactory coding in a diverse group of terrestrial animals.

## Methods

### List of Species

#### Nematocera

##### Culicidae

*Aedes aegypti* (L.) *Aedes albopictus* (Skuse), *Anopehels albimanus* (Wiedemann), *Anopheles albimanus* (Wiedemann), *Anopheles arabiensis* (Patton), *Anopheles atroparvus* (Van Thiel), *Anopheles bwambae* (White), *Anopheles christyi* (Newstead & Carter), *Anopheles coluzzii* (Coetzee & Wikerson), *Anopheles culicifacies* (Giles), *Anopheles darlingi* (Root), *Anopheles dirus* (Peyton & Harrison), *Anopheles epiroticus* (Linton & Harbach), *Anopheles farauti* (Laveran), *Anopheles funestus* (Giles), *Anopheles gambiae* (Giles), *Anopheles maculatus* (Theobald), *Anopheles melas* (Theobald), *Anopheles merus* (Dönitz), *Anopheles minimus* (Theobald), *Anopheles quadriannulatus* (Theobald)*, Anopheles sinensis* (Wiedemann), *Anopheles stephensi* (Liston), *Culex pipiens quinquefasicatus* (L.)

##### Psychodidae

*Phlebotomus papatasi* (Scopoli), *Lutzomyia longipalpis* (Lutz & Neiva)

#### Brachycera

##### Muscidae

*Musca domestica* (L.), *Stomoxys calcitrans* (L.), *Haematobia irritans* (L.)

##### Glossinidae

*Glossina austeni* (Newstead), *Glossina brevipalpis* (Newstead), *Glossina fuscipes* (Newstead), *Glossina morsitans* (Westwood), *Glossina pallidipes* (Austen), *Glossina palpalis* (Robineau-Desvoidy)

##### Drosophilidae

*Drosophila melanogaster* (Meigen)

### Identification of IndolORs and Gene Annotations

*IndolOR* homologs were identified in the genomes if Dipteran species listed above via tBLASTn or BLASTp searches against available genome assemblies on the National Center for Biotechnology Information (www.ncbi.nlm.nih.gov) or Vectorbase (www.vectorbase.org). Gene annotations were corrected based upon multiple amino acid alignments using Geneious Prime^2019^ software (Biomatters Limited, USA), as well as conservation of intron positions. Chromosomal synteny was determined by identifying conserved orthologs encoded in either the 5’ or 3’ directions of indolORs on their respective genome assemblies.

### Phylogenetic Analysis

Conceptual amino acid sequences of 92 *indolORs* were aligned using MAFFT version 7 [26]. The evolutionary history was inferred by using the Maximum Likelihood method and JTT matrix-based model [27]. The tree with the highest log likelihood (−20016.97) is shown. The percentage of trees in which the associated taxa clustered together is shown next to the branches. Initial tree(s) for the heuristic search were obtained automatically by applying Neighbor-Join and BioNJ algorithms to a matrix of pairwise distances estimated using the JTT model, and then selecting the topology with superior log likelihood value. The tree is drawn to scale, with branch lengths measured in the number of substitutions per site. This analysis involved 92 amino acid sequences with a total of 431 positions in the final dataset. Evolutionary analyses were conducted in MEGA X [28, 29]. One thousand bootstrap pseudoreplicates were performed and best tree was obtained, where branches with >50% bootstrap were retained.

### Chemical reagents

The chemicals used for the deorphanization of MdomOR30a were obtained from Acros Organics (Morris, NJ), Alfa Aesar (Ward Hill, MA), and Thermo Fisher Scientific (Waltham, MA) at the highest purity available. Compounds were initially dissolved to 1M in 100% DMSO to generate stock solutions and serially diluted in ND96 perfusion buffer (96mM NaCl, 2mM KCl, 5mM MgCl2, 0.8mM CaCl2, and 5mM HEPES) for oocyte recordings.

### Two-electrode voltage clamp of Xenopus laevis oocytes

MdomOrco and MdomOR30a coding regions were obtained by *de novo* synthesis into a pENTR plasmid from a commercial source (Twist Bioscience, South San Francisco, CA). Subcloning into the pSP64t expression plasmid was accomplished using the Gateway^®^ LR clonase II^®^ enzyme (Life Technologies, Carlsbad, CA). Individual cRNAs were synthesized *in vitro* from linearized pSP64t using the mMESSAGE mMACHINE^®^ SP6 kit (Life Technologies). Stage V-VII *Xenopus laevis* oocytes were obtained from a commercial source (Xenopus1, Dexter, MI) and incubated in ND96 incubation media (96 mM NaCl, 2mM KCl, 5mM HEPES, 1.8mM CaCl_2_, 1mM MgCl_2_, pH 7.6) supplemented with 5% dialyzed horse serum, 50 μg/mL tetracycline, 100μg/mL streptomycin, 100μg/mL penicillin, and 550 μg/mL sodium pyruvate. Oocytes were injected with 27.6 nL (27.6 ng of each cRNA) of RNA using a Nanoliter 2010 injector (World Precision Instruments, Inc., Sarasota, FL). Odorant-induced currents of oocytes expressing AgamOrco and MdomOR30a were recorded using the two-microelectrode voltage-clamp technique (TEVC) using an OC-725C oocyte clamp (Warner Instruments, LLC, Hamden, CT), while maintaining a holding potential of −80 mV. Data acquisition and analysis were carried out with the Digidata 1550B digitizer and pCLAMP10 software (Molecular Devices, Sunnyvale, CA, USA). A panel of 12 structurally-related indolic compounds: Indole-3-acetic acid, 2,3-dioxoindoline, 4-hydroxyindole, quinoline, indole-3-carboxyaldehyde, 2,3-dimethylindole, 3-methylindole, indole, methyl indole-3-carboxylate, 2-methylindole, 2-oxindole, and methyl salicylate, were used to determine the most efficacious ligand for MdomOR30a (n=14 oocytes). Compounds were perfused at 10^−4^ M for 10 s and current was allowed to return to baseline between administrations. Kurtosis was calculated for the raw amplitudes and the KURT function in Microsoft Excel 365. Concentration-dependent responses were determined by perfusing oocytes with indole, 3-methylindole, or 2,3-dimethylindole across a log-molar range of 10^−9^ to 10^−3^ M. Data analyses were performed using GraphPad Prism 8 (GraphPad Software Inc., La Jolla, CA, USA). Compounds were perfused for up to 30s or until peak amplitude was reached. Current was allowed to return to baseline between chemical compound administrations.

## Supporting information

Amino acid sequence of fly indolORs

Odorant concentrations used in TEVC of Xenopus oocytes expressing MdomOR30a

Gene structure of fly indolOR

Genomic locations of fly indolOR genes

## Supplementary information

Supplementary information accompanies this paper at: **Additional file1: Tables S1–S4. Table S1:** List of indolORs, abbreviations, and amino acid sequences by species. **Table S2:** Raw oocyte recording data. **Table S3:** IndolOR gene annotations. **Table S4:** Nematocera and brachycera syntenic relationships.

## Acknowledgements

We acknowledge Robert Huff (Baylor University) for assistance with electrophysiology and Amir Dekel (Hebrew University of Jerusalem) for assistance with kurtosis calculation.

## Authors’ contributions

RJP and JDB conceived and designed the study. RJP and JDB conducted the bioinformatics analyses. SJS performed electrophysiology experiments. RJP and JDB analyzed the data and wrote the manuscript. All authors read and approved the final manuscript.

## Availability of data and materials

The datasets supporting the conclusions of this article are included within the article and its additional files.

## Ethics approval and consent to participate

Not applicable.

## Author details

1 Department of Biology, Baylor University, Waco, Texas, United States of America

2 Department of Entomology, The Hebrew University of Jerusalem, Rehovot, 76100, Israel

